# Synaptogenic activity of the axon guidance molecule Robo2 is critical for hippocampal circuit function

**DOI:** 10.1101/840710

**Authors:** Heike Blockus, Sebastian V. Rolotti, Miklos Szoboszlay, Tiffany Ming, Anna Schroeder, Kristel M. Vennekens, Phinikoula Katsamba, Fabiana Bahna, Seetha Mannepalli, Goran Ahlsen, Barry Honig, Lawrence Shapiro, Joris de Wit, Attila Losonczy, Franck Polleux

## Abstract

The developmental transition between axon guidance and synapse formation is critical for circuit assembly but still poorly understood at the molecular level. We hypothesized that this key transition could be regulated by axon guidance cues switching their function to regulate synaptogenesis with subcellular specificity. Here, we report evidence for such a functional switch, describing a novel role for the axon guidance molecule Robo2 in excitatory synapse formation onto dendrites of CA1 pyramidal neurons (PNs) in the mouse hippocampus. Cell-autonomous deletion of Robo2 from CA1 PNs leads to a drastic reduction of the number of excitatory synapses specifically in proximal dendritic compartments. At the molecular level, we show that this novel postsynaptic function of Robo2 depends on both its canonical ligand Slit and a novel interaction with presynaptic Neurexins. Biophysical analysis reveals that Robo2 binds directly to Neurexins via its Ig4-5 domains. *In vivo* 2-photon Ca^2+^ imaging of CA1 PNs during spatial navigation in mice shows that sparse deletion of Robo2 during development drastically reduces the likelihood of place cell emergence and alters spatial coding properties of the remaining place cells. Our results identify Robo2 as a novel molecular effector linking synaptic specificity to the acquisition of spatial coding properties characterizing hippocampal circuits.

---

Proper circuit function relies on the establishment of synaptic connections characterized by a high degree of cell type and subcellular specificity. How this striking degree of synaptic specificity is achieved during development remains poorly understood, especially in the complex brains of mammals. Many cell surface molecules that mediate molecular recognition between axons and dendrites during circuit development have been identified^1–4^. These trans-synaptic adhesion molecules are often expressed in a cell-type specific manner^5^ thereby determining the pool of a cell’s possible synaptic partners. Some classes of these molecules have synaptic organizing properties evidenced by their direct or indirect ability to recruit key pre- and postsynaptic proteins. When axons reach their target areas, a cellular switch from a phase of growth and branching to synaptogenesis is observed. This switch is characterized by dynamic instability of adhesion progressively leading to more stable patterns of synaptic connectivity characterizing adult circuits. One of the foremost outstanding questions in neuroscience is how synapse formation is mechanistically integrated with earlier developmental steps such as axon guidance and branching. One potential mechanism for coordinating this transition relies on the possibility that axon guidance cues acquire synaptogenic functions during circuit wiring. Indeed, evidence for such dual use of developmental molecules has been reported^6^. However, it remains unclear what prompts molecules to switch their function from axon guidance to synaptogenesis. Here we report that the well-studied axon guidance ligand-receptor pair Slit/Robo has a hitherto unknown function in synaptogenesis that depends on a novel interaction with the presynaptic organizing transmembrane proteins Neurexin at nascent synapses. Slit-Robo signaling has been extensively studied for its role in axon guidance^7^ and branching ^8^ for over three decades across many model organisms, but a possible role for Slit-Robo signaling in synaptogenesis and circuit function remains largely unexplored. Among all axon guidance receptor/ligand pairs, we focused on Robo/Slit to test their potential role in synaptic specificity for two main reasons: (1) their expression pattern in the hippocampus and cortex is maintained at stages of circuit development following completion of axon guidance: Robo/Slit expression spans phases of synaptogenesis into adulthood strongly suggesting that Robo/Slit signaling could be involved in steps of neuronal development beyond axon guidance; (2) recent work demonstrated that a large class of transmembrane proteins regulating synaptic specificity contain Leucine-rich repeat (LRR) domains^9^, and interestingly, the extracellular secreted ligands Slit1-3 contains four LRR domains.

Our results reveal a Slit-dependent postsynaptic function for Robo2 in instructing excitatory, but not inhibitory, synapse formation via a trans-synaptic adhesion complex with the presynaptic organizing molecules Neurexins. We show that Robo2 is critical for the establishment of the synaptic architecture characterizing hippocampal CA1 PNs *in vivo*, which play a key role in navigation, episodic learning and memory. Excitatory glutamatergic inputs are anatomically segregated within the dendritic arbor of CA1 PNs: inputs from intrahippocampal CA3 and CA2 regions arrive onto proximal dendrites, while distal apical dendrites are primarily innervated by long-range inputs from the entorhinal cortex (EC)^10^ (Figure 1e). It is known that information conveyed by these inputs streams is qualitatively distinct, and that the temporal integration of these inputs governs *in vivo* response properties of CA1 PNs – most notably the encoding of the animal’s position in a given environment by a subset of spatially tuned cells (termed ‘place cells’)^11^. Hence, CA1 PNs are an ideal model to relate molecular mechanisms of synapse-specific input compartmentalization to the emergence of *in vivo* physiological properties. Strikingly, we found that the Robo2 protein is expressed in an input-specific manner - spatially restricted to basal and proximal apical dendrites of CA1 PNs. Using 2-photon calcium imaging in awake behaving mice, we demonstrate that cell-autonomous deletion of Robo2 from CA1 PNs reduces their probability of becoming place cells, and the Robo2-deficient place cells that do remain are functionally different from their wild-type (WT) counterparts. Altogether, our results identify a novel function for Slit-Robo signaling, beyond axon guidance, in the mediation of synaptic specificity through formation of a trans-synaptic complex with presynaptic Neurexins. We demonstrate Robo2’s impact on excitatory synapse formation through its cell-autonomous deletion from CA1 PNs, which alters their spatial coding properties in the hippocampus. Our results provide a missing link between the molecular mechanisms underlying synaptic specificity and the emergence of circuit properties during mammalian brain development.

**Figure 1.**
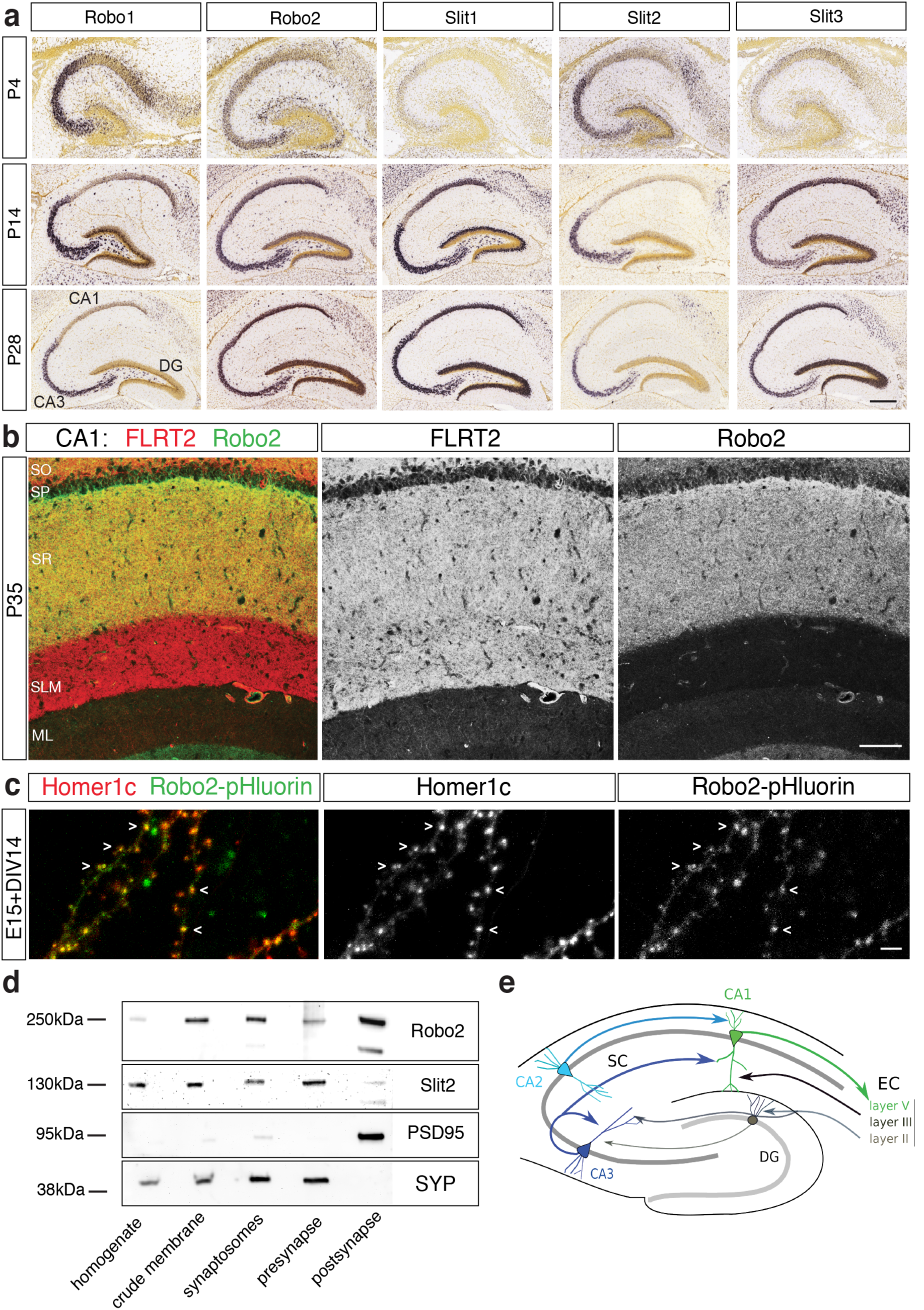
Robo2 expression pattern in the hippocampus and its subcellular localization at synapses. **a.** Publicly available mRNA expression pattern of Robo1, Robo2 and Slit1,2,3 in the early postnatal and mature mouse hippocampus. Robo2, Slit1,3 are expressed throughout CA regions in the hippocampus, Robo1 and Slit2 expression is confined to CA3. Robo2 expression increases between P4 and P14. Scalebar: 500µm. **b.** Immunohistochemistry for FLRT2 (red) and Robo2 (green) in the rat hippocampus at P35. While FLRT2 is expressed throughout CA1, expression of Robo2 is confined to SO and SR. Scalebar: 100µm. SR, stratum radiatum; SLM, stratum lacunosum-moleculare; ML, molecular layer of DG; SO, stratum oriens; SP, stratum pyramidale. **c.** Subcellular localization of Robo2 *in vitro. Ex utero* electroporation of Robo2-pHluorin and Homer1c-tdtomato in pyramidal neurons *in vitro*. Robo2-pHluorin colocalizes with the excitatory postsynaptic marker Homer1c at the majority of dendritic spines at 14 DIV. Scale bar: 15µm. **d.** Synaptic fractionation and immunoblot for Robo2, Slit2 and synaptic markers. Successful isolation of pre-and postsynaptic compartments evidenced by presence of Synaptophysin and PSD95 respectively. Robo2 localizes to pre-and postsynaptic compartments, its ligand Slit2 is enriched presynaptically. **e.** Schematic of hippocampal circuitry.

## RESULTS

### Robo2 is expressed in the developing and mature hippocampus and localizes to excitatory synapses

To determine whether Slit/Robo could be involved in stages of neuronal development after completion of axon guidance, we first characterized its region-specific and subcellular localization in the postnatal mouse hippocampus. Using publicly accessible resources (Allen Brain Atlas), we found that while almost absent at P4, Robo2 mRNA is expressed throughout all Cornu Ammonis (CA) regions of the hippocampus at P14, and its expression is maintained into adulthood (P56, Figure 1a). Expression of Robo1 and the Robo ligand Slit2 within the hippocampus is confined mostly to the CA3 region, which provides presynaptic input to CA1. Since Robo2 but not Robo1 is expressed in CA1, this allowed us to isolate functions of Robo2 without redundancy to Robo1 therein. We next determined Robo2 protein expression at P35 in CA1. Strikingly, Robo2 protein is expressed specifically in CA1 stratum oriens (SO) and stratum radiatum (SR), but is not detected in stratum lacunosum moleculare (SLM) (Figure 1b,e). SO and SR correspond to layers where axons from CA3 and CA2 PNs synapse onto dendrites CA1 PNs, while the apical tuft of CA1 PNs in SLM receives long-range inputs from the entorhinal cortex (EC). To gain more insights into the postsynaptic localization of Robo2 at excitatory synapses received by PNs, we used *ex utero* electroporation to express a pHluorin-tagged version of Robo2 in pyramidal neurons *in vitro*^12^. A pHluorin tag fused to the extracellular domain of Robo2 allows the specific visualization of the plasma membrane targeted form of Robo2, but not the pool of the protein contained in intracellular vesicles^13^. This approach shows that Robo2-pHluorin co-localizes with the excitatory postsynaptic marker Homer1c in spines of pyramidal neurons in culture (E15+14 days in vitro (DIV)) corresponding to the peak of synaptogenesis *in vitro* (Figure 1c). To further delineate which synaptic compartment Robo and Slit localize to, we performed biochemical synaptic fractionation. Using western blotting, we found that Robo2 is enriched in postsynaptic membranes and that its ligand Slit2 is enriched in presynaptic membranes (Figure 1d).

### Robo2 is required for excitatory synapse formation in CA1 pyramidal neurons *in vivo*

Having established that Robo2 localizes to synapses and enriched postsynaptically, we next determined whether Robo2 was required cell-autonomously for excitatory synaptic development in CA1 PNs *in vivo*. To accomplish this, we used *in utero* electroporation (IUE) to cell-autonomously delete Robo2 from a subset of CA1 PNs using a conditional Robo2 knockout mouse (Robo2^F/F^). We performed hippocampal IUE (HIUE) of Cre-recombinase expressing plasmid together with a non-limiting amount of a plasmid encoding a Cre-dependent reporter (flex-tdTomato) into CA1 progenitors of control (wild-type) and Robo2^F/F^ embryos at embryonic day E15.5 (Figure 2a). This approach is not only cell-autonomous with regard to CA1 PNs, but effectively postsynaptic-autonomous because the axons of CA1 PNs form almost no recurrent excitatory connections with other CA1 PNs^14^ and therfore almost all presynaptic inputs to the electroporated CA1 PNs are wild-type. Using this approach, we found that conditional deletion of Robo2 from CA1 PNs throughout development leads to a significant (∼40%) reduction of spine density in proximal dendritic compartments (SO: basal, 39.67%, SR: apical oblique 41.98%), but does not affect spine density in distal apical tuft dendrites (SLM: tuft) (Figure 2b). Dendritic growth of Robo2-deficient CA1 PNs was not affected (Figure S1a-c). Taken together with its protein subcellular localization pattern (Figure 1b), our results demonstrate that Robo2 is required postsynaptically for excitatory synaptic development in an input-specific manner in CA1 PNs.

**Figure 2.**
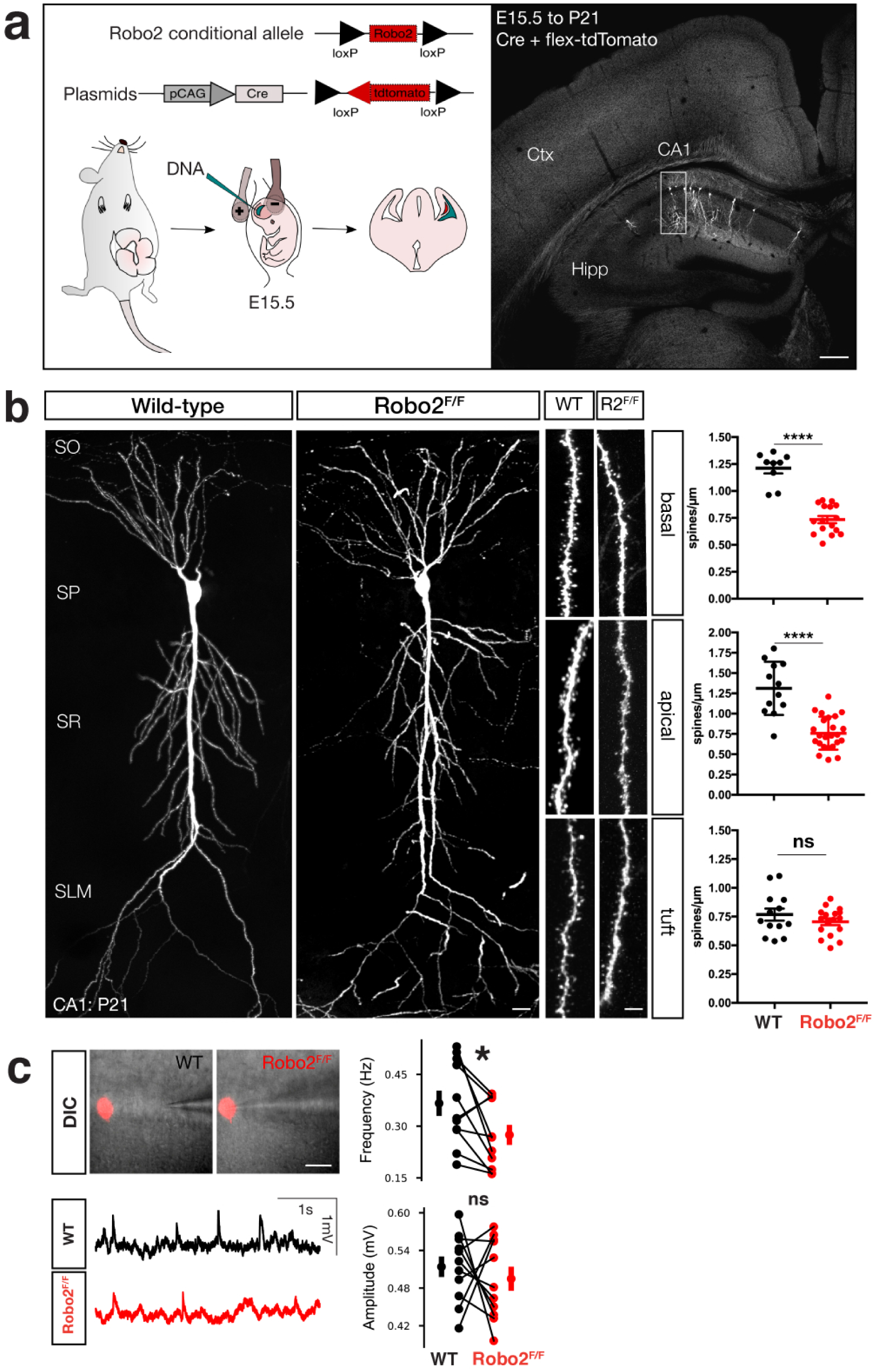
Robo2 is required for synapse formation in hippocampal CA1 PNs. **a.** Schematic of *in utero* electroporation to sparsely target CA1 PN progenitors in order to introduce plasmids expressing Cre-recombinase and flex-tdtomato. **b.** Optically isolated CA1 PNs in either WT or Robo2^F/F^ mice. Analysis of spine architecture reveals a decrease in spine density in proximal dendritic compartments of Robo2-deficient CA1 PNs (basal: WT: n=9 dendritic segments, mean=1.21 spines/µm +/-0.049 (SEM), KO: n=16 dendritic segments, mean=0.73 spines/µm +/- 0.033 (SEM), reduction=39.67%, ****p<0.0001, Mann-Whitney; apical: WT: n=12 dendritic segments, mean=1.31 spines/µm +/-0.095 (SEM), KO: n=24 dendritic segments, mean=0.76 spines/µm +/-0.041 (SEM), reduction=41.98%, ****p<0.0001, Mann-Whitney), but not distal tuft dendrites (tuft: WT: n=13 dendritic segments, mean=0.76 spines/µm +/-0.052 (SEM), KO: n=17 dendritic segments, mean=0.71 spines/µm +/-0.029 (SEM), p=0.51, ns, Mann-Whitney). Scalebar: 25µm. **c.** Whole-cell patch clamp and sEPSP measurements of WT and Robo2-deficient CA1 PNs. DIC images and example traces of patched Robo2 KO neuron as indicated by presence of flex-mcherry and neighboring WT neuron (n=11 pairs). Quantification of amplitude (mV, WT avg: 0.514; SD: 0.052 mV, KO avg: 0.495; SD: 0.061 mV, p=0.54592, paired t-test, ns), area under the curve (mV*ms, WT avg: 16.512; SD: 3.363 mV*ms, KO avg: 14.975; SD: 2.912 mV*ms, p=0.39907, paired t-test, ns), inter-event interval (IEI, s, WT avg: 3050.710; SD: 1061.686 ms, KO avg: 4094.111; SD: 1389.291 ms, *p=0.01297, paired t-test) and frequency (Hz, WT avg: 0.366; SD: 0.116 Hz, KO avg: 0.275; SD: 0.092 Hz, *p=0.0329, Wilcoxon-signed rank test).

To confirm that our sparse, conditional, deletion of Robo2 affected the synaptic physiology of CA1 PNs, we turned to patch-clamp recordings in acute adult hippocampal slices. Using the same HIUE approach in Robo2^F/F^ mice to achieve a sparse, mosaic deletion of Robo2 from CA1 PNs, we performed whole-cell current-clamp recordings of spontaneous excitatory postsynaptic potentials (sEPSP) in Robo2 KO (tdTomato-expressing) and neighboring WT CA1 PNs within the same hippocampal slices (Figure 2c and Figure S1b). As expected from the reduction in spine density, the frequency of sEPSPs was significantly reduced in Robo2-null compared to adjacent WT CA1 PNs, whereas their amplitude was not significantly different (Figure 2c). These data demonstrate that postsynaptic Robo2 expression is required for excitatory synapse development in CA1 PNs.

### Robo2 induces excitatory synapse formation in a Slit-dependent manner

The data described thus far show that Robo2 localizes to and is required for excitatory synapse development in CA1 PNs in a compartment-specific manner. Next, we used a reductionist experimental setting to determine whether Robo is directly involved in synapse formation. Specifically, we used an *in vitro* hemisynapse assay^15^ to determine whether Robo proteins expressed on the surface of HEK293 cells could induce formation of presynaptic boutons from axons of co-cultured primary neurons. Using Neuroligin1 (NLG1) as a positive control since it can induce both excitatory and inhibitory synapses^15^ and CD8 as negative control, we tested whether Robo receptors were able to induce the formation of presynaptic boutons from the axons of co-cultured cortical neurons. Indeed, expression of Robo1 and Robo2 in HEK293 cells led to a strong clustering of axonal Vglut1 around the cell perimeter (Figure 3a). Interestingly, Robo3, did not induce accumulation of Vglut1+ presynaptic boutons. Robo3 is a divergent member of the Robo family and has lost the ability to bind Slit ligands during mammalian evolution^16^. The finding that Robo3 is not able to induce Vglut1+ presynaptic bouton clustering prompted us to determine if the synaptogenic activity of Robo was Slit-dependent. To do so, we expressed a Robo2-receptor with a deletion of the Slit binding domain (Robo2^ΔIg1,2, 17^) in the HEK293 cells and found that Robo2^ΔIg1,2^ did not induce Vglut1+ presynaptic bouton clustering. Some synaptogenic transmembrane proteins such as NLG1 indiscriminately induce the formation of excitatory and inhibitory presynaptic boutons in the hemisynapse assay^15^. By contrast, HEK cells expressing Robo2 did not induce clustering of the presynaptic inhibitory Vgat1 from surrounding axons (Figure 3b). Altogether, our data show that Robo1/2 specifically induces the formation of excitatory, but not inhibitory, synapses in a Slit-dependent manner.

**Figure 3.**
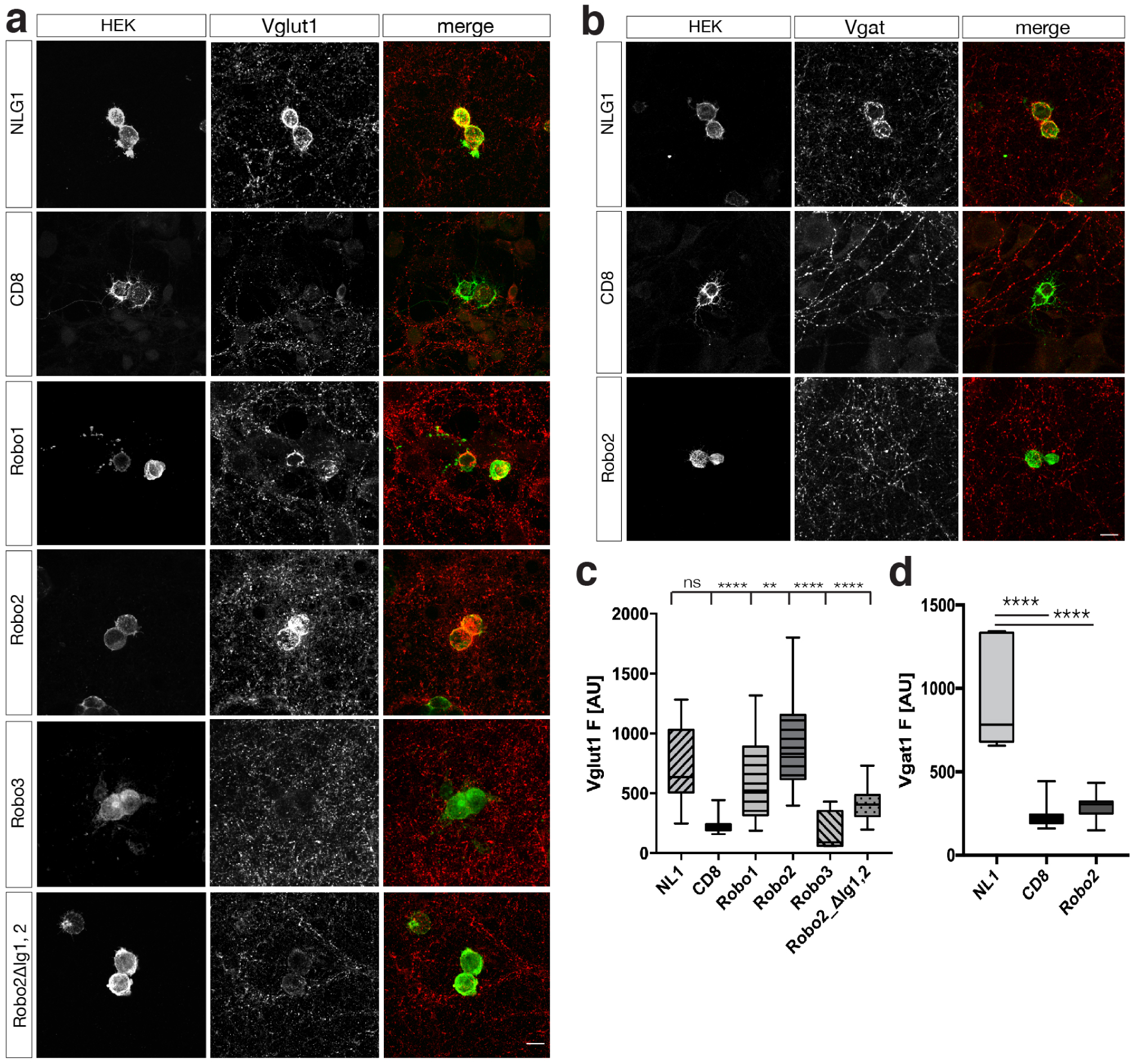
Robo2 induces excitatory synapse formation in a Slit-dependent manner. **a.** *In vitro* hemisynapse assay (24h coculture of primary neurons at DIV12 with HEK293 cells expressing cDNAs as indicated in text boxes on the left for each row of images). Immunostaining for Vglut1 indicates presence of excitatory presynaptic boutons, which cluster around the perimeter of the HEK293 cell in case the cDNA is synaptogenic. NLG1: positive control, CD8: negative control. Robo1 and Robo2 expression in HEK293 cells leads to Vglut1 clustering around the cell perimeter. Both Slit-binding deficient receptors, Robo3 and Robo2^ΔIg1,2^, are unable to induce Vglut1 clustering. Scale bar: 7µm. **b.** Robo2 does not induce inhibitory synapse formation as evidence by the absence of clustering the inhibitory presynaptic marker Vgat1 in immunohistochemistry. Scale bar: 7µm. **c,d.** Quantification of (A,B): one-way ANOVA, ****p<0.0001, ***p<0.001, **p<0.01, *p<0.05, whiskers show min/max, n=4 independent experiments).

### Presynaptic Neurexins bind directly to Robo2 and are required for Robo2-dependent synaptogenesis

Our *in vitro* and *in vivo* results show that Robo2 is both necessary and sufficient to induce excitatory synapses, prompting us to determine the identity of the trans-synaptic binding partners on the presynaptic membrane required for Robo2’s synaptogenic activity. We hypothesized that Robo2 may be part of a trans-synaptic adhesion complex with its secreted ligand Slit and an unknown presynaptic partner. Since (1) homophilic trans-interactions of Robos can occur in vivo at least in the context of axon guidance in *Drosophila*^18^, (2) axonal function of Robo as an axon guidance receptor is well known and (3) we detected a small fraction of the Robo2 pool on presynaptic membranes (Figure 1d), we first sought to test whether the synaptogenic activity of Robo1/2 requires the presence of Robo1/2 expressed in axons/presynaptically. To address this, we repeated the hemi-synapse assay using co-culture with cortical neurons from Robo1 knockout or Robo1/2 double knockout embryos. Since a homozygous constitutive deletion of Robo2 (but not Robo1) is perinatally lethal^19^, we isolated cortical neurons from a combination of Robo1 constitutive knockout and our Robo2 conditional allele and infected the cultures with Cre-lentivirus at DIV0 (Figure 4a). Interestingly, Robo1/2-deficient axons were still able to cluster Vglut1 around HEK293 cells expressing Robo2. Our results show that presynaptic Robo1/2 expression is not required to support the synaptogenic function of postsynaptic Robo2. We therefore hypothesized that a novel presynaptic transmembrane protein meditating the synaptogenic function of Robo-Slit. In order to identify the interactome of Slit and Robo at synapses, we took an unbiased proteomic approach based on an experimental pipeline developed recently^20^. We purified synaptosome fractions from P21 rat brains and used it to perform a pulldown with recombinant Slit2-Fc protein (Supplementary Figure 2a). Shotgun mass-spectrometry (MS) analysis of synaptic proteins binding to recombinant Slit2-Fc identified Neurexin1/2/3 as one of the transmembrane proteins pulled down by Slit2. We also identified Robo2, Glypican1 and PlexinA1, surface proteins previously identified as Slit-interacting proteins^21–23^, validating our approach. We focused on Neurexins as potential presynaptic Slit-Robo interacting partners mediating their synaptogenic activity because of the well-characterized function of Neurexins as a presynaptic organizing protein family^2^.

**Figure 4.**
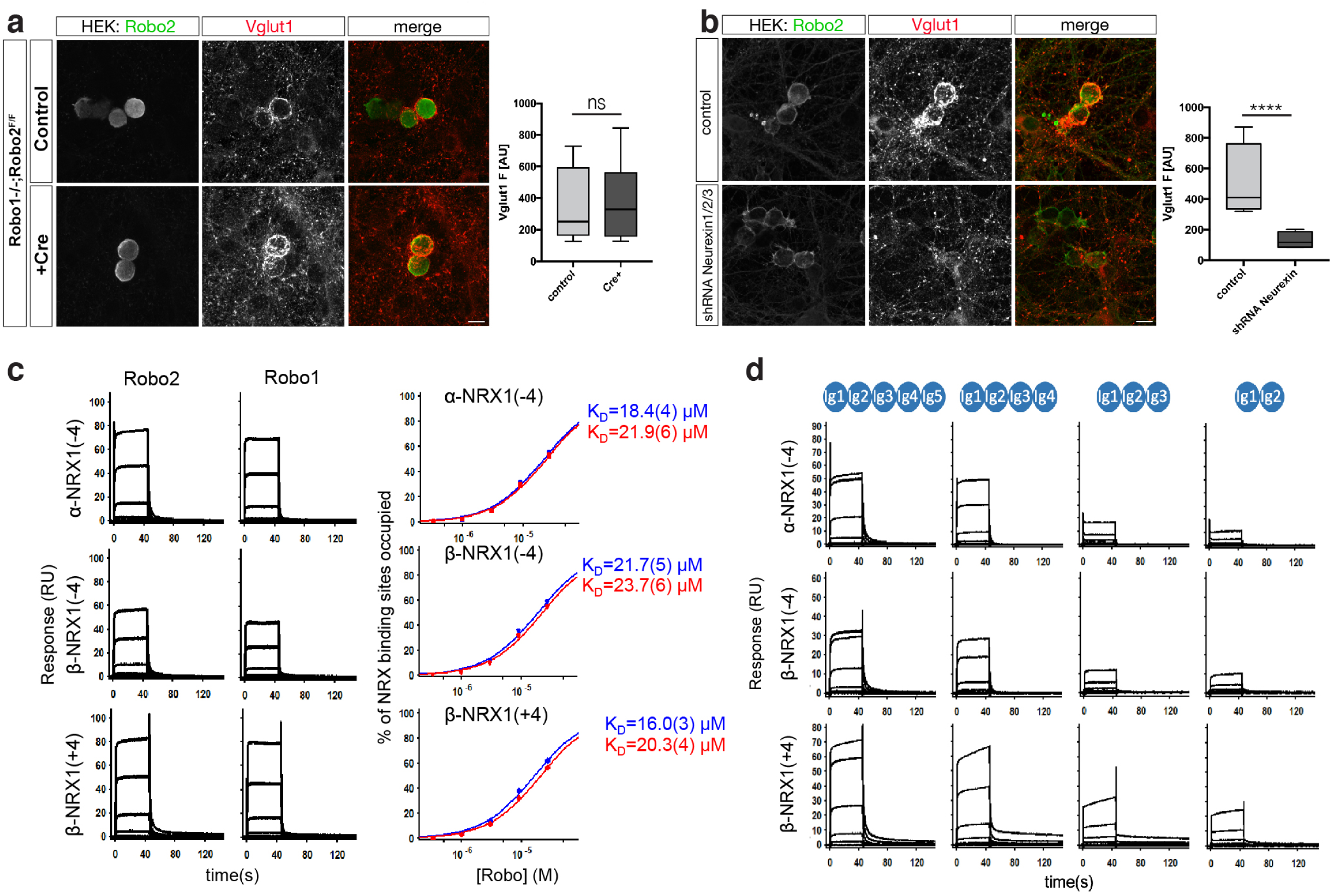
Robo2 is part of a heterophilic complex with presynaptic Neurexins in *trans*. **a.** Presynaptic Robo1/2 are not required for Vglut1 clustering around Robo2-expressing HEK293 cells. *In vitro* hemisynapse assay was performed using primary neurons from either Robo1^-/-^ or Robo1^-/-^;Robo2^F/F^ (infected with Lentivirus expressing Cre-recombinase at DIV0, one-way ANOVA, ****p<0.0001, ***p<0.001, **p<0.01, *p<0.05, whiskers show min/max, n=3 independent experiments). Scale bar: 7µm. **b.** Presynaptic Neurexins are essential for Robo2-dependent Vglut1-clustering. *In vitro* hemisynapse assay was performed in the presence (DIV3) of a Lentivirus expressing a pan-Neurexin shRNA. Neurexin knockdown completely abolishes Robo2-dependent Vglut1-clustering. Scale bar: 7µm. **c.** SPR binding experiments of Robo1 and Robo2 extodomains over NRX surfaces: Binding of Robo2 and Robo1 (Ig1-5) over surfaces immobilized with α-NRX1Δ4, β-NRX1Δ4, and β-NRX1+4 ectodomains. Robo 2 and Robo1-binding was detected with all three NRX-immobilized surfaces. Binding isotherms of the percentage of NRX binding sites occupied vs Robo concentration (right panels) yield the K_D_s for Robo2 (Ig1-5) and Robo1 (Ig1-5), shown in blue and red, respectively. The number in brackets represents the error of the fit. **d.** Binding of Robo2 Ig deletion fragments over α-NRX1Δ4, β-NRX1Δ4, and β-NRX1+4 ectodomains. Robo2 fragments encompassing Ig domains 1-5, 1-4, 1-3 and 1-2 respectively were tested for binding. A sharp decrease in signal is observed between the Ig1-4 and Ig1-3 fragments.

In order to determine whether presynaptic Neurexins are required for the synaptogenic function of Robo2, we repeated the hemi-synapse assay after knockdown of presynaptic Neurexins^24^. Neurexins are key presynaptic organizing proteins capable of forming multiple trans-synaptic complexes with postsynaptic proteins in a synapse-specific way. Three main Neurexin genes (*Nrx1-3*) can each generate two main isoforms (α (long) and β (short)) that display a considerable degree of alternative splicing. As a first approach, we used shRNA targeting all isoforms of Neurexin1/2/3α and β and infected primary neurons before plating the HEK293 cells expressing Robo2 for the hemi-synapse assay. Strikingly, Neurexin1/2/3(α/β)-knockdown completely abolished Robo2-synaptogenic activity (Figure 4b), establishing presynaptic Neurexins as an essential component of the trans-synaptic adhesion complex mediating Slit-dependent Robo2 synaptogenic activity.

To further characterize the potential interaction between Neurexins, Slit and Robo2 at synapses, we turned to surface plasmon resonance (SPR), a biophysical method to analyze direct interactions between macromolecules allowing quantitative measurement of binding affinities. Given the high functional diversity of different Neurexin isoforms, we chose to immobilize three Neurexin1 isoforms: α-Neurexin1 without insertion of the major splice-site (SS) 4 (α-NRX1^(−4)^), β-Neurexin without or with SS4 (β-NRX1^(−4)^ and β-NRX1^(+4)^ respectively) on a dextran-chip via amide coupling and flowed purified recombinant Robo1 and Robo2 (Ig domains 1-5; Supplementary Figure 2f) over the chip as analyte (Figure 4c). Robo1 and Robo2 bound directly to all Neurexin isoforms tested with K_D_s of ∼16-23µM, an affinity in the same order of magnitude as β-Neurexin1/2/3-Neuroligin1/2/3 interactions (ranges between ∼0.8-56µM)^25^. Interestingly, we did not observe any direct interaction of Slits and Neurexins in SPR (data not shown). However, the fact that we identified Robo in our Slit2-Fc pull-down/MS experiments from synaptosomes (Supplementary Figure 2a) suggest that Nrxn identification in the same experiment was due to its interaction with Robo.

To start identifying the binding interface between Neurexins and Robo1/2, we performed similar SPR experiments with Ig domain deletions of Robo2. We found a drastic reduction in binding efficiency of Robo2 to Neurexins when Ig4-5 were deleted (Figure 4d). This reveals that binding sites for Slit (Ig1-2) and Neurexins (Ig4-5) on Robo2 do not overlap. Notably, Robo-Neurexin interactions were dependent on the presence of Heparin as well as calcium (Supplementary Figure 2b-d). Together with our findings that the Slit-binding domain is important for Robo2-dependent synaptogenesis (Figure 3a) these data suggest the existence of a tripartite Robo-Slit-Neurexin complex promoting excitatory synapse formation.

### Sparse developmental deletion of Robo2 alters place cell properties of CA1 PNs *in vivo*

We then sought to determine whether interfering with Robo2-dependent development of excitatory synapses in CA1 PNs affects their coding properties *in vivo*. CA1 PNs represent an ideal model to study the impact of interfering with synaptic specificity on circuit function. A subset of CA1 PNs exhibit spatially tuned firing when the animal is exploring an environment^11^. The emergence of place coding is in large part governed by the convergence of spatially tuned excitatory inputs from upstream hippocampal regions CA2/CA3 onto CA1 PNs ^26, 27^. We assessed CA1 PN place cell properties at the population level using *in vivo* two-photon (2p) microscopy-based Ca^2+^ imaging in head-fixed, awake behaving mice^28–30^.

We used the same HIUE approach to conditionally delete Robo2 via Cre expression alongside Cre-dependent mCherry from a sparse subpopulation of CA1 PNs in Robo2^F/F^ animals. This allows direct comparison of response properties in WT and Robo2 KO cells within the same animal (Figure 5a). We then broadly infected dorsal CA1 via stereotactic injection with a recombinant adeno-associated virus (rAAV) expressing the genetically-encoded Ca^2+^ indicator GCaMP6f. This allows us to image the activity of hundreds of GCaMP6f-expressing CA1 PNs within stratum pyramidale amongst which ∼10% of all imaged neurons are deleted for Robo2 as identified by the presence of Cre-dependent mCherry expression (Figure 5b-d). Mice were trained to run for randomly delivered water rewards (random foraging) on a linear treadmill belt decorated with spatial cues as navigational landmarks.

**Figure 5.**
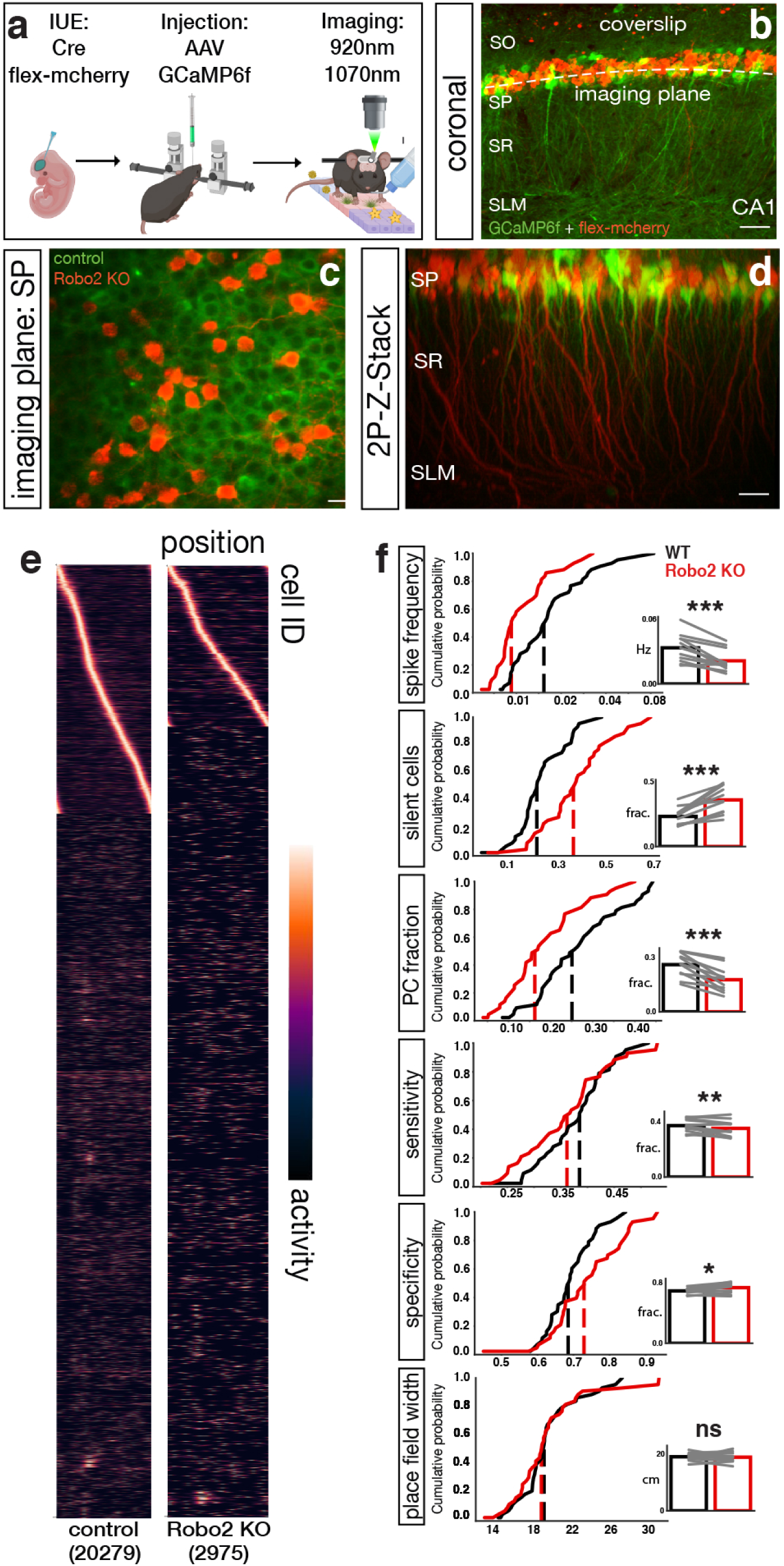
Robo2 knockout alters place cell properties of CA1 PNs *in vivo*. **a.** Overview of the imaging paradigm. Once adult, mice were injected with AAV-hSyn-GCaMP6f and implanted with a metal headpost to allow for *in vivo* 2-photon imaging in awake-mice. Mice explore a treadmill belt with spatial cues in a 1D environment. **b.** Representative image of field of view (*post-hoc*). HIUE was used to create a mosaic Robo2 knockout of CA1 PN subpopulations. Scale bar top left: 35µm. **c.** Imaging field of view acquired during a session. Scale bar 15µm. **d.** *In vivo* Z-stack volume of imaged area. Scale bar bottom right: 20µm. **e.** Normalized tuning of all imaged cells (total cell counts in parentheses; place cells sorted by peak) across sessions for WT and Robo2 KO neurons. **f.** Differences in spike frequency and place cell properties between WT and Robo2 KO neurons. Robo2 KO neurons exhibit a decrease in spike frequency, place cell fraction and sensitivity, but a modest increase in specificity. Details see text (paired t-test; spike frequency (Hz): ***p=0.0004, fraction silent cells: ***p=0.0003, sensitivity: **p= 0.01, specificity: *p= 0.05, fraction place cells ****p=0.0001, place field width (cm): p=0.6, ns; n= 11 FOVs) CDFs show distribution of means across imaging sessions. Dashed lines indicate median. *insets*: pairwise comparisons after aggregation across FOV sessions.

We then analyzed inferred spike rates from deconvolved Ca^2+^ activity and compared place cell properties of Robo2 KO cells and their WT counterparts in adult mice (>60 days). We found that Robo2 KO CA1 PNs have greatly reduced spiking frequency during running (WT: 0.033±0.013 Hz, KO: 0.022±0.010 Hz; mean±sd) (Figure 5f). Overall, we observed a significant (∼40%) reduction in the fraction of place cells within Robo2 KO compared to WT CA1 PNs (WT: 0.260±0.066, KO: 0.177±0.069) (Figure 5e-f). Furthermore, the remaining Robo2 KO place cells showed significant alterations in their response properties. Relative to WT CA1 PNs, Robo2 KO CA1 PNs showed significant reduction in their sensitivity (WT: 0.373±0.048, KO: 0.353±0.058), defined as the reliability of spiking during place field traversals. Interestingly, we observed a slight but significant increase in specificity of spiking activity in the Robo2 KO compared to WT CA1 PNs (WT: 0.686±0.038, KO: 0.730±0.067), suggesting that the decrease in Robo2 KO spike frequency has an outsize effect on out-of-field firing. No difference in place field width was observed (WT: 18.87±1.39 cm, KO: 18.73±1.72). In sum, our data shows that Robo2-dependent alteration in excitatory synapse development has a significant impact on *in vivo* coding properties of hippocampal CA1 PNs in awake behaving mice.

## DISCUSSION

We have uncovered a novel role for the axon guidance molecules Slit and Robo in excitatory synapse development in CA1 pyramidal neurons of the hippocampus. Our data demonstrate postsynaptic Robo promotes excitatory (but not inhibitory) synapse formation by forming a trans-synaptic complex with its LRR domain-containing Slit ligand and presynaptic transmembrane Neurexins. Interestingly, Robo2 protein localization is restricted to specific dendritic domains of CA1 PNs (apical oblique and basal dendrites but not apical tufts), and conditional deletion of Robo2 leads to a selective ∼40% decrease in excitatory synapse formation within these two domains corresponding to dendritic domains receiving excitatory inputs from CA2/CA3 PNs. Finally, we found that cell-autonomous deletion of Robo2 from CA1 PNs alters their place cell properties *in vivo*. Altogether, our data shows for the first time that Robo2 functions outside of axon guidance in excitatory synapse formation and therefore plays a vital role in the emergence of CA1 PNs coding properties and hippocampal circuit function (Figure 6).

**Figure 6.**
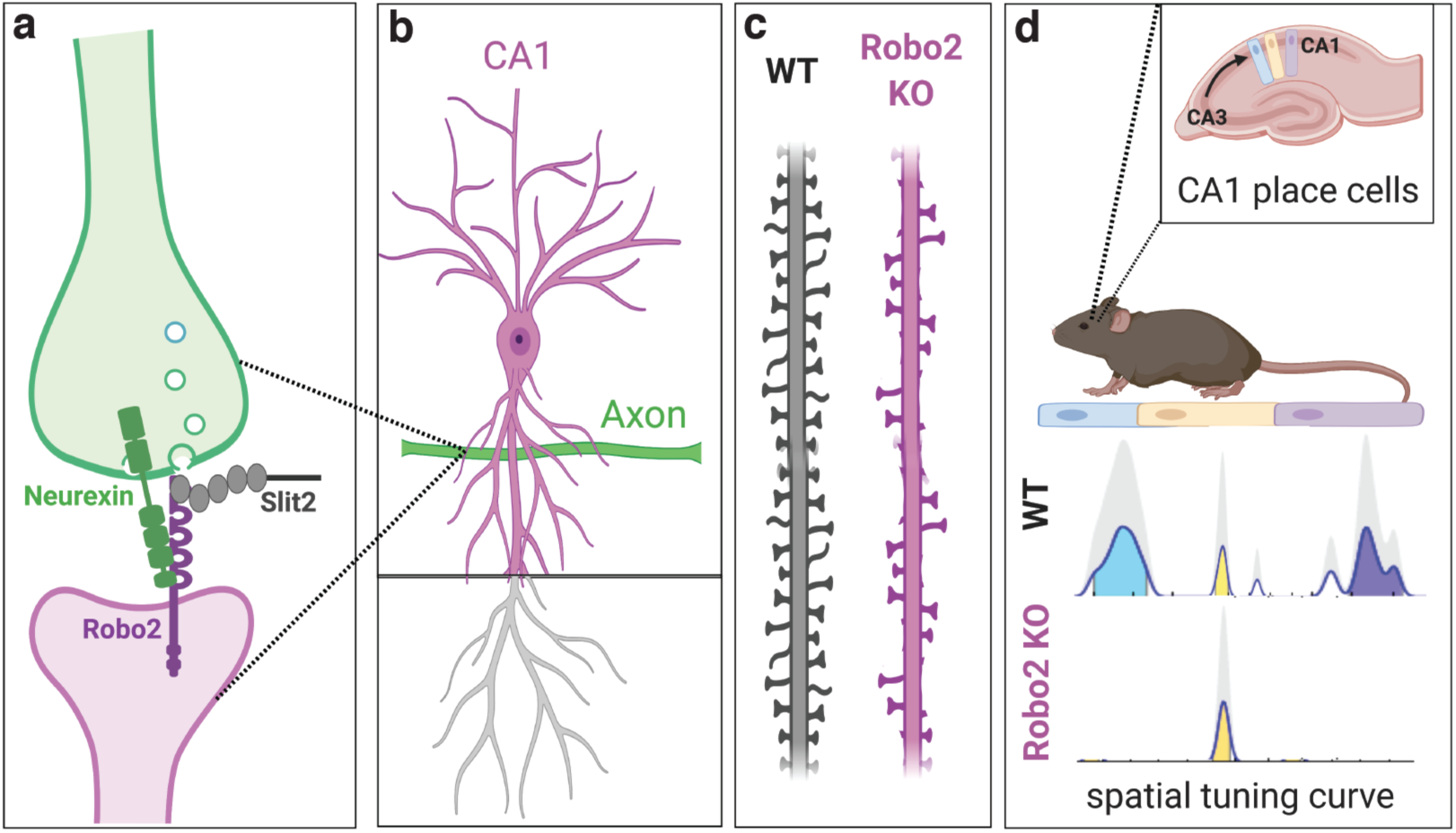
Schematic summary of main conclusions. **a.** Robo2’s synaptogenic function relies on Slit and presynaptic Neurexins, to which Robos bind directly *in vitro*. **b.** Robo2 is expressed in proximal dendritic compartments of CA1 PNs. **c.** In line with the expression pattern, deletion of Robo2 from CA1 PNs leads to a decrease in spine density in basal and proximal apical compartments. **d.** Cell-autonomous deletion of Robo2 from CA1 PNs reduces the fraction of CA1 neurons that are spatially tuned, but neuronal activity in Robo2-deficient CA1 PNs is slightly, but significantly more specific to its place field location.

### Slit-Robo signaling in axon guidance and synapse formation

In the context of axon guidance, overwhelming evidence has demonstrated that Slit binding to Robo elicits signaling leading to growth cone repulsion^31^. Interestingly, a study has suggested that in dendrites of cortical pyramidal neurons Slit-Robo signaling elicits a chemoattractive response and also promotes dendritic branching^32^. Our study reveals that in the context of synapse formation, postsynaptic Robo promotes excitatory synapse formation in a Slit-and Neurexin-dependent manner. Together with our biochemical data, the most parsimonious model we can propose is that during the switch between axon guidance and synapse formation, Slit binding to axonal Robo receptors elicits chemorepulsion but that during synapse formation, dendritic Robo receptors elicit excitatory synapse formation by forming a trans-synaptic complex with Slit and presynaptic Neurexins. Slit binds through its LRR domains to the first two Ig domains (Ig1-2) of Robo^17^ and our results show that Neurexins bind most efficiently to Ig4-5 of Robo. Furthermore, recent structural work shows that the Robo trans-dimerization domain is contained within Ig4-5^33^. We therefore propose a model whereby Slit-binding to Robo releases the inhibitory Robo trans-interaction inducing a conformational change^33^ which might allow postsynaptic Robo to bind to presynaptic Neurexins. However, using recombinant proteins in our SPR assay, α-and β-Neurexin1 can bind to Robo in the absence of Slit but it is possible that in these conditions Robo is rather unconstrained conformationally and therefore does not require Slit to enable Neurexin-Robo interaction.

Furthermore, the source of Slit in Robo2-dependent synaptogenesis remains an open question in the hippocampus since Slit2 expression is restricted CA3 i.e. presynaptic to CA1 PNs but Slit1 and Slit3 are detected both in CA1 and CA3. We speculate that Slit2 expressed by incoming CA3 axons reaching CA1 PN dendrites is the relevant source of Slit for the synaptogenic function of postsynaptic Robo2 in CA1 PNs. In future experiments, it will be important to improve our understanding of the structural mechanisms underlying the Robo-Slit-Neurexin trans-synaptic complex to further dissect how these proteins interact in a context-specific manner.

### Trans-synaptic coincidence detection as a molecular mechanism to increase synaptic specificity

Our results uncover a novel trans-synaptic molecular complex constituted of Robo and Neurexin. The finding that Slit as well as Neurexin are both important to support Robo2-dependent synaptogenesis *in* vitro, posits the existence of a tripartite trans-synaptic protein complex. This would suggest a model whereby synaptic specificity during recognition of a presynaptic axon and the corresponding postsynaptic dendrites requires coincidence detection at the molecular level between at least three components as recently exemplified^34^. We identified other transmembrane components in our synaptic Slit pull-down mass spectrometry experiments such as Glypican1 and PlexinA1 (previously characterized Slit binding proteins^22, 23^) which could further increase the specificity of this trans-synaptic molecular complex by decreasing the probability that each component would be present at axon-dendrite contacts during synapse formation. We also reveal that Neurexin-Robo interaction requires heparin-conjugated moieties and that the synaptogenic activity of Robo is abolished by treatment with heparinase. Since Neurexin1 was recently shown to require a rare glycan modification, heparan sulfate^35^, this posttranslational modification might also participate in increasing the specificity of trans-synaptic interactions between Robo, Slit and Neurexins. Future experiments will need to provide insights into the structure of this and other trans-synaptic protein complex in order to gain further evidence for this coincidence detection model as a mechanism to increase synaptic specificity.

### Input-specific reduction of excitatory drive alters place cell properties

Our results show that cell-autonomous deletion of Robo2 from CA1 PNs reduces the number of inputs they receive from CA2/CA3 onto their basal and apical oblique dendrites by ∼40% (see detailed numbers above), which has significant consequences for their ability to encode spatial information. Hence, our molecular manipulation serves as a tool to precisely reduce the impact of intrahippocampal information on CA1 PNs, while leaving long-range cortical inputs intact. This allows us to change the quantity of these different inputs onto the CA1 PN output function. CA3/CA2 provides spatially tuned input to CA1 PNs and we find that if these inputs are reduced, Robo2-deficient CA1 PNs are less sensitive for their given place field. Our results emphasize the striking ability of CA1 PNs to compartmentalize their proteome within their dendritic arbor, which translates not only to compartmentalized inputs along different dendritic domains but also ultimately leads to their unique physiological properties^36–39^.

Altogether, our study demonstrates how the precise localization and interactome of developmentally relevant proteins determine the functional properties of neurons within mature circuits through their ability to regulate synaptic specificity and circuit connectivity.

## Acknowledgments

We thank Miyako Hirabayashi and Qiaolian Liu for technical assistance and Alain Chédotal for providing Robo2 conditional and Robo1 constitutive knockout mice. We thank members of the Polleux and Losonczy labs for stimulating discussions. This work was supported by NIH-NINDS (RO1NS067557) (FP), NINDS (R21NS109753-01A1) (FP, AL, HB) and an award from the Roger De Spoelberch Foundation (FP). AS was supported by FWO PhD fellowship (11Z3715N/17N); JdW is supported by FWO Odysseus grant, FWO project grant G0C4518N, and FWO EOS grant G0H2818N. Mass spectrometry analysis was performed at the VIB Proteomics Core (Leuven, Belgium). AL is further supported by the National Institute of Mental Health (NIMH) 1R01MH100631, National Institute of Neurological Disorders and Stroke (NINDS) 1U19NS104590 and 1R01NS094668, the Zegar Family Foundation Award, and the Kavli Foundation. B.H. is supported by a National Science Foundation grant (MCB-1412472), L.S. by NIH grant R01MH114817.

## Author contributions

H.B. and F.P. designed the study and wrote the manuscript. H.B. performed hemisynapse assays, primary culture overexpression experiments, hippocampal *in utero* electroporation and structural analysis of Robo2 KO neurons (aided by T.M.), as well as *in vivo* 2-photon calcium imaging experiments. S.V.R. analyzed calcium imaging data. A.L. oversaw 2-photon calcium imaging experiments and analysis. M.S. performed *in vitro* electrophysiological experiments and analyzed the data. A.S. performed immunohistochemistry for Robo2. K.V. performed synaptic fractionation. J.dW. oversaw and designed experiments performed by A.S and K.V. and helped with interpretation of some of the experiments. F.B., S.M. and G.A. performed all recombinant protein production and purification. P.K. performed SPR experiments. L.S. and B.H. oversaw and designed experiments performed by F.B., S.M. and G.A..

## Materials & Correspondence

All correspondence and requests for material should be addressed to.

## Methods

### Animals

Mice were used according to protocols approved by the Institutional Animal Care and Use Committee (IACUC) at Columbia University and in accordance with National Institutes of Health guidelines. The health and welfare of the animals was supervised by a designated veterinarian. The Columbia University animal facilities comply with all appropriate standards (cages, space per animal, temperature, light, humidity, food, water). Both males and females were used for all experiments. To the best of our knowledge, we are not aware of an influence of sex on the parameters analyzed in this study. Timed-pregnant *CD1* females were purchased from Charles Rivers. *129/SvJ, C57Bl/6J* nontransgenic mice and Robo1^-/-^;Robo2^F/F^ and Robo2^F/F^ transgenic mice were maintained in a 12-hour light/dark cycle. Timed-pregnant females were obtained by overnight breeding with males of the same strain. Noon the day after the breeding was considered as E0.5.

### Lentivirus production

Second generation VSV.G pseudotyped lentiviruses were produced as previously described^40, 41^. HEK293T cells were transfected with control or shRNA-containing pFUGW (GFP-expressing) vector plasmids and helper plasmids PAX2 and VSVG using Fugene (Promega). Supernatant was collected 65 hr after transfection, spun at 2000 rpm to remove debris and filtered through a 0.45 μm filter (Corning). In order to maximize the purity of viral particle pellets, a small amount of sucrose solution (20% sucrose, 100 mM NaCl, 20 mM HEPES, 1 mM EDTA, at pH 7.4, filtered with a 0.22 μm filter) was placed at the bottom of the centrifuge tubes before adding filtered media. Viral particles were then pelleted by centrifugation at 200.000 rpm for 2 hr at 4C. Final pellet was re-suspended in 200 μl of PBS and stored at −80°C in 10 μl aliquots.

### Immunohistochemistry

Immunofluorescent staining for synaptic proteins in hippocampal sections was performed in 16 μm-thick cryosections from P35 rat brains perfused with 2% paraformaldehyde (PFA) solution. Sections were post-fixed in a 1:1 MeOH:acetone solution at −20°C, and then permeabilized with 0.5% triton in PBS-0.2% gelatin. Blocking solution consisted of 10% normal horse serum and 0.5% triton in PBS-0.2% gelatin. Primary antibodies were the following: goat anti-FLRT2 (R&D Systems) and rabbit anti-Robo2 (Aviva Systems). Primary and secondary antibodies were diluted in 5% normal horse serum and 0.5% triton in PBS-0.2% gelatin. Hoechst was used as a nuclear stain (5nM in PBS). Prolong Gold Antifade (ThermoScientific) was used to mount slides. Confocal images were taken on a Leica TCS SP5 microscope or on a Zeiss LSM880 confocal with an Airyscan detector.

### *Ex utero* electroporation and primary neuron culture

Cortices from E15.5 mouse embryos were dissected followed by dissociation in complete Hank’s balanced salt solution (cHBSS) containing papain (Worthington) and DNase I (100ug/mL, Sigma) for 15 minutes at 37°C, washed three times, and manually triturated in DNase I (100ug/mL) containing neurobasal medium (Life Technology) supplemented with B27 (1x, Thermo Fischer Scientific,), FBS (2.5%, Gibson) N2 (1x, Thermo Fischer Scientific), glutaMAX (2mM, Gibco). Cells were plated at 10^5 cells per 12 mm glass coverslip pre-coated with Poly-D-Lysine and Laminin (Corning). One-third of the medium was changed every 7 days thereafter with non-FBS containing medium and maintained for 11-21 days in 5% CO2 incubator at 37°C. *Ex utero* electroporation was performed as previously described^42^. Plasmids used for *ex utero* electroporation were all in pCAG vector backbone^43^ expressing the following cDNAs: pCAG-Robo2-pHluorin^12^, Homer1c-tdtomato.

### Synaptic fractionation and Western Blotting

Synaptic fractionation was based on a previously described method^44^. Briefly, 20 P21 mice brains were homogenized in 10 mL per 2.5 brains with Solution A (0.32 M Sucrose-1 mM NaHC0_3_, 1mM MgCl_2_, 0.5 mM Cacl_2_ and protease inhibitors) (homogenate), centrifuged at 1,500 x g for 15 min, and the supernatant was collected (post nuclear supernatant). The supernatant was then centrifuged at 14,000 x g for 20 min, and the resulting supernatant (cytosol) and pellet (crude membrane) collected. The pellet was re-suspended in 24ml Solution B (0.32 M sucrose-1mM NaHC0_3_ and protease inhibitors) and loaded onto 0.5 M/1.0 M/1.2 M discontinuous sucrose gradients and centrifuged at 32500 x g for 120 min. The material at the 1.0 M/ 1.2 M interface was collected (synaptosome). The fraction was diluted with Solution B to 60ml. Triton X-100 was added to 0.5% and extracted at 4°C by stirring for 15 min. The extract was centrifuged at 32,500 x g for 25 min, the supernatant collected (soluble synaptosome/Triton-soluble fraction) and the pellet was re-suspended in Solution B, loaded onto a 1.0 M/1.5 M/2.0 M sucrose gradient, and centrifuged at 200,000 x g for 2 hr. Material was collected at the 1.5M/2.0M interface (PSD). 0.5% Triton X-100 was added and detergent-soluble material extracted at 4°C by end-over-end agitation for 10 min. Lastly, the extract was centrifuged at 200,000 x g for 20 min and the pellet re-suspended in Buffer B +5% SDS (purified PSD/Triton-insoluble fraction).

Samples were then prepared for western blotting in Laemmli buffer heated at 60C for 10mins and 20µg were run on a 4-20% Acrylamide gel (Biorad.cat no 456-1093). Wet western blotting was carried out using 0.2µm Nitrocellulose membranes (GE Healthcare Amersham, Protean cat no 10600015). Membranes were blocked in TBS-T, 5% Milk and incubated with the following primary antibodies overnight at 4C: mouse anti-PSD-95 (ThermoScientific), rabbit anti-c-Myc (Santa Cruz Biotechnology), rabbit anti-HA (Sigma), rabbit anti-βIII-tubulin (Abcam), mouse anti-Synaptophysin (Sigma), mouse anti-β-actin (Sigma). The following day, blots were washed in TBS-T and incubated with HRP coupled secondary antibodies according to the manufacturer’s instructions. Signals were revealed using ECL Super signal West Femto (ThermoScientific cat no 34095).

### *In utero* hippocampal electroporation and spine analysis

*In utero* electroporation targeting the hippocampus was performed using a triple-electrode setup as previously described^45^ to target hippocampal CA1 PN progenitors at E15.5. Plasmids were injected into the lateral ventricle (pCAG-Cre: 0.3µg/µl, pEFIA-tdTomato/mCherry 0.5µg/µl) followed by 5 pulses at 45V (50ms duration, 500ms interpulse interval). Mice were perfused transcardially at P21 and sectioned at 200µm on a vibratome (Leica). Sections were mounted and imaged on a Nikon W1 spinning disk microscope using a 100x, 1.45 NA silicon-immersion objective. Spine analysis was carried out using vaa3d (detection parameters as in Iascone et al 2019, https://www.biorxiv.org/content/10.1101/395384v1) by three independent experimenters blinded to experimental condition.

### *In vitro* electrophysiology

#### Slice preparation

Mice of either sex (total of n=5, n=2 female, n=3 male, between P24-P27) were anesthetized with isoflurane, decapitated and the brain was placed in ice cold dissection solution containing (in mM): 195 sucrose, 10 NaCl, 15 glucose, 26 NaHCO_3_, 2.5 KCl, 1.25 NaH_2_PO_4_, 1 CaCl_2_*2H_2_O, 2 MgCl_2_*6H_2_0. 400 µm-thick horizontal slices were cut using a Leica VT1200 vibratome. After dissection, slices were incubated in an ACSF-containing submerged chamber at 32°C for 30 minutes, then stored at room temperature until used for recording. ASCF contained the following (in mM): 125 NaCl, 10 glucose, 26 NaHCO_3_, 2 Na-pyruvate, 2.5 KCl, 1.25 NaH_2_PO_4_, 2 CaCl_2_*2H_2_O, 1 MgCl_2_*6H_2_0. Brain slices were used until up to 6h post-dissection. Both dissection and ACSF solutions were carbonated to saturation, resulting in a pH of 7.3-7.4 and osmolality of 310 ± 5 mOsm.

#### Whole-cell recordings

Whole-cell current clamp recordings were performed using a K-gluconate-based internal solution (in mM: 130 K-gluconate, 8 KCl, 10 HEPES, 4 NaCl, 4 Mg_2_ATP, 0.3 Tris_2_GTP, 14 Tris-phosphocreatine and 6 biocytin with pH = 7.28 and osmolality 295 mOsm) at 32°C in ACSF. Thick-walled borosilicate glass pipettes (outer diam.: 1.5 mm, inner diam.: 0.86 mm; Sutter Instruments) were pulled with a P-97 pipette puller (Sutter Instruments). Open tip resistance when filled with internal solution was 1.7-4.7 MΩ (average: 2.9 ± 0.8 MΩ, n=22 cells). Data were acquired with using a MultiClamp 700B amplifier (Molecular Devices) and a CED Micro1401-3 digitizer (Cambridge Electronic Design Limited). Data were digitized at 50 kHz and low-pass filtered at 1 kHz. Series resistance and pipette capacitance were monitored continuously and compensated throughout the 10-minute recording sessions. Data were discarded when series resistance exceeded 20 MΩ (average: 9.6 ± 4.7 MΩ, n=22 cells) and the holding current decreased below −100 pA for −60 mV. After sequentially patching one KO (identified by mCherry fluorescence, Supplementary Figure 1d-f) and a neighboring WT CA1 pyramidal cell, the brain slices were fixed in 4% PFA solution for *post hoc* histological processing. Slices were incubated overnight with Streptavidin-Alexa488 (Thermo Fisher) diluted 1/1000 in 4% normal goat serum, 0.2% TritonX-100 in PBS. EPSPs were detected by a template matching algorithm and analyzed in Stimfit^46^.

### *In vitro* hemisynapse assay and immunocytochemistry

Mixed culture assays were performed as previously described^47^. Briefly, HEK293T cells were transfected with cDNAs for NLG1, CD8, Robo1, Robo2, Robo3, Robo2ΔIg1,2 using Fugene6 (Promega), mechanically dissociated and cocultured with 11 DIV cortical neurons for 8-24 hr, depending on the experiment. Following coculture, the cells were immunostained for Vglut1, Vgat1, GFP, myc and HA. For the analysis of heparinase III treatment, neurons (11 DIV) were treated with 1 U/ml heparinase III (Sigma-Aldrich) or vehicle (20 mM Tris-HCl [pH 7.5], 0.1 mg/ml BSA, 4mM CaCl_2_) for 2 hr at 37°C. Cells were then washed twice with neuronal feeding medium and subsequently cocultured with transfected HEK293T cells for an additional 8 hr. Cells were then processed for immunostaining. Coverslips were washed once with 500uL of DPBS and fixed with 4%PFA, 1X PBS, 4% sucrose solution at RT for 15 minutes. Cells were washed with PBS 3X for 5 minutes and permeabilized in 0.2% TritonX-100 in 1xPBS for 15mins. Cells were blocked in 1%BSA, 4%NGS for 30mins at RT. Primary incubation was done overnight at 4C: HA (abcam, sheep), GFP (Aves, chicken), Vglut1 (SySy, guinea-pig), Vgat1 (SySy, mouse), Myc (Thermo Fisher, mouse). The following day, cells were washed in 1xPBS 3x for 5mins followed by incubation with secondary antibodies (Invitrogen-Alexa-dye-coupled) for 1h at RT. Cells were then washed in 1xPBS 3x for 5mins and mounted in Fluoromount-G (Southern Biotech cat no 0100-01). Images were then acquired using a NikonA1R confocal microscope using a 60X, 1.4 NA oil-immersion objective. To test the effect of neuronal Robo1/2 knockout, neurons from Robo1^-/-^;Robo2^F/F^ mice were infected with lentivirus expressing Cre-recombinase under a synapsin promoter at DIV0. To test the effect of neuronal Neurexin knockdown, neurons were infected with lentivirus expressing a previously validated shRNA against Neurexins^24^ at DIV3. For quantification of mixed-culture assays, images were thresholded using ImageJ and the total area of Vglut1 puncta was measured and normalized to the total GFP-positive area per cell. Individual puncta could not be measured in this assay because thresholding resulted in the fusion of individual puncta due to their high density. Measurements were performed in a minimum of three independent preparations and in each experiment for any given condition a minimum of twenty cells were acquired. Imaging and analysis were conducted blind to the condition.

### Protein expression and purification for SPR

We used cDNAs encoding rat Robo1 (NCBI: NP 071524.1), rat Robo2 (NCBI: NP 115289.1), and rat α-neurexin (NCBI: NP 068535.2) as templates for PCR amplification of desired coding regions for expression constructs. Each protein was produced into the pLEXm mammalian cell expression vector proceeded by a BiP signal peptide and in frame with a C-terminal octahistidine tag. The Robo1 construct encoded amino acids for Ig 1-5 (Gly58-Phe545). For Robo2, constructs and corresponding reside ranges were Ig1-2 (Gly21-Glu226), Ig1-3 (Gly21-Ala315), Ig1-4 (Gly21-Asp413) and Ig1-5 (Gly21-Ser509). The α-neurexin construct lacked the splice insertion sequences at splice sites 1, 2 and 4 (Δ1Δ2Δ4) and encoded amino acids (Leu31-Thr1336). β-neurexin1 Δ4 encoded Thr1125-Val1327, and β-neurexin1+4 encoded Thr1125-Val1327 (amino acid numbering based on NCBI: NP 068535.2). All proteins were expressed in Human Embryonic Kidney (HEK) 293 Freestyle cells (Invitrogen) in suspension culture using serum-free media. Plasmid constructs were transfected into HEK293 cells using polyethyleneimine (Polysciences). Cell growths were harvested five days after transfection, and the secreted proteins were purified from supernatant using nickel affinity chromatography followed by size exclusion chromatography in either 10 mM Tris pH 8.0 and150 mM sodium chloride for Robo1/2 or 10 mM Tris pH 8.0, 150 mM sodium chloride and 3 mM calcium chloride for neurexins.

### SPR binding experiments

SPR binding experiments were performed using a Biacore T100 biosensor equipped with a Series S CM4 sensor chip. α-Neurexin1^(−4)^, β-Neurexin1^(−4)^, β-Neurexin1^(+4)^ were immobilized over individual flow cells using amine-coupling chemistry in HBS-P pH 7.4 (10mM HEPES-OH, 150mM NaCl, 0.005% Tween-20) buffer supplemented with 3mM CaCl_2_, at 32°C using a flow rate of 20μL/min. Prior to immobilization, the three Neurexin protein samples were buffer exchanged in HBS pH 7.4, 3mM CaCl_2_ using Zeba spin desalting columns (Thermo Scientific) prior to immobilization. Dextran surfaces were activated for 7 minutes using equal volumes of 0.1 M NHS(N-Hydroxysuccinimide) and 0.4 M EDC(1-Ethyl-3-(3-dimethylaminopropyl)carbodiimide). Each protein of interest was immobilized at ∼40μg/mL in 10 mM sodium acetate, pH 5.5, 3mM CaCl_2_ until the desired immobilization level was achieved. The immobilized surface was blocked using a 3-minute injection of 1.0 M ethanolamine, pH 8.5. Each molecule was immobilized at 1600-1700 RU. An unmodified surface was used as a reference flow cell to subtract bulk shift refractive index changes.

Binding experiments were performed at 25°C in a running buffer containing 10 mM Tris-Cl pH 8.0, 150 mM NaCl, 3mM CaCl_2_, 10 μg/mL heparin, 1 mg/mL BSA and 0.005% (v/v) Tween-20. Binding analysis in the absence of heparin was tested in the same buffer lacking heparin and experiments without CaCl_2_ were performed in a buffer containing 3mM EGTA instead of CaCl_2_. Robo2 and 1 (Ig1-5) and Robo2 Ig fragment analytes were prepared in running buffer using a three-fold dilution series at 27, 9, 3, 1 and 0.333μM, except for the EGTA experiment, where binding was tested at the highest concentration of 27 μM. In each binding cycle, analytes were injected over all immobilized surfaces at 50μL/min for 45s, followed by 180s of dissociation phase, a running buffer wash step and a buffer injection at 100μL/min for 60s. Each series was tested in order of increasing concentration and then repeated in the same experiment to confirm the reproducibility of the binding assay. Running buffer was used instead of an analyte every two cycles, to double reference the binding responses by removing systematic noise and instrument drift. The binding signal between 40 and 44 seconds for each analyte, was fit against the Robo concentration using a 1:1 interaction model to calculate the K_D_. The data was processed and analyzed using Scrubber 2.0 (BioLogic Software).

### Fc-Protein Purification for Mass-spectrometry

Fc protein purification was performed as described previously^20^. Slit2-Fc (N-(aa1-840) and C-terminal (aa841-1290) fragments) proteins were produced by transient transfection of HEK293T cells using PEI (Polysciences). Six hours after transfection, media was changed to OptiMEM (Invitrogen) and harvested 5 days later. Conditioned media was centrifuged, sterile-filtered and run over a fast-flow Protein-G agarose (Thermo-Fisher) column. After extensive washing with wash buffer (50 mM Hepes pH 7.4, 300 mM NaCl and protease inhibitors), the column was eluted with Pierce elution buffer. Eluted fractions containing proteins were pooled and dialyzed with PBS using a Slide-A-Lyzer (Pierce) and concentrated using Amicon Ultra centrifugal units (Millipore). The integrity and purity of the purified ecto-Fc proteins was confirmed with SDS-PAGE and Coomassie staining, and concentration was determined using a Bradford protein assay.

### Affinity Chromatography for Mass-spectrometry

Affinity chromatography experiments were performed as previously described^20^. Crude synaptosome extracts were prepared from ten P21-22 rat brains, homogenized in homogenization buffer (4 mM Hepes pH 7.4, 0.32 M sucrose and protease inhibitors) using a Dounce homogenizer. Homogenate was spun at 1,000 x g for 10 minutes at 4°C. Supernatant was spun at 14,000 x g for 20 minutes at 4°C. P2 crude synaptosomes were re-suspended in Extraction Buffer (50 mm Hepes pH 7.4, 0.1 M NaCl, 2 mM CaCl_2_, 2.5 mM MgCl_2_ and protease inhibitors), extracted with 1% Triton X-100 for 2 hours and centrifuged at 100,000 x g for 1 hour at 4°C to pellet insoluble material. Fast-flow Protein-A sepharose beads (GE Healthcare) (250µl slurry) pre-bound in Extraction Buffer to 100 µg human Fc or Slit2-Fc were added to the supernatant and rotated overnight at 4°C.

Beads were packed into Poly-prep chromatography columns (BioRad) and washed with 50 ml of high-salt wash buffer (50 mM HEPES pH 7.4, 300 mM NaCl, 0.1 mM CaCl2, 5% glycerol and protease inhibitors), followed by a wash with 10 ml low-salt wash buffer (50 mM HEPES pH 7.4, 150 mM NaCl, 0.1 mM CaCl2, 5% glycerol and protease inhibitors). Bound proteins were eluted from the beads by incubation with Pierce elution buffer and TCA-precipitated overnight. The precipitate was re-suspended in 8 M Urea with ProteaseMax (Promega) per the manufacturer’s instruction. The samples were subsequently reduced by 20-minute incubation with 5mM TCEP0 (tris(2carboxyethyl)phosphine) at RT and alkylated in the dark by treatment with 10 mM Iodoacetamide for 20 additional minutes. The proteins were digested overnight at 37°C with Sequencing Grade Modified Trypsin (Promega) and the reaction was stopped by acidification. Mass spectrometry analysis was performed by the VIB Proteomics Core (Leuven, Belgium).

### Stereotactic virus injection and craniotomy

For imaging experiments, recombinant adeno-associated viruses carrying the GCaMP6f gene were obtained from the Penn Vector Core (AAV1.Syn.GCaMP6f.WPRE.SV40) with titer of 2– 4 × 10^13^. Dorsal CA1 was stereotactically injected at −2.0 mm AP; −1.5 mm ML; and −0.9, −1.2, −1.4 mm DV relative to the cortical surface. Mice were then surgically implanted with an imaging window over the left dorsal CA1 and implanted with a stainless-steel headpost for head fixation during imaging experiments.

### *In vivo* two-photon calcium imaging

We used the same imaging system as described previously ^28, 48^. All images were acquired with a Nikon 40× NIR water-immersion objective (0.8 NA, 3.5 mm WD) in distilled water. For excitation, we used a Chameleon, Ultra II (Coherent) laser tuned to 920 nm, and a Fidelity-2 (Coherent) laser at 1070 nm. We continuously acquired red (mCherry) and green (GCaMP6f) channels separated by an emission cube set (green, HQ525/70 m-2p; red, HQ607/45 m-2p; 575dcxr, Chroma Technology) at 512 × 512 pixels covering 330 μm × 330 μm at 30 Hz with photomultiplier tubes (green GCaMP6f fluorescence, GaAsP PMT, Hamamatsu Model 7422P-40; red mCherry fluorescence, GaAsP PMT Hamamatsu). For four mice, red and green channels were recorded simultaneously with both lasers simultaneously exciting. For three mice, only the green channel was excited/recorded during behavior, while a red-only image for cell identification was acquired immediately prior to recording. A custom dual stage preamp (1.4 × 10^5^ dB, Bruker) was used to amplify signals prior to digitization.

For training, mice were water restricted (>90% pre-deprivation weight) and trained to run on a cue-poor burlap treadmill belt for a non-operantly delivered water reward over the course of 1– 2 weeks. We applied a progressively restrictive water reward schedule, with mice initially receiving 12 randomly placed reward zones per lap and ultimately receiving 1 randomly placed reward zone per lap. Mice were habituated to the optical instrumentation, then trained for 20 min daily until they regularly ran at least one lap per minute. During imaging sessions, mice received one randomly placed water reward per lap, with water delivered for every subsequent lick inside the reward zone for a maximum of 2.5 s. The reward zone position was changed randomly each lap.

### Imaging Analysis

#### Processing of Ca^2+^ Fluorescence Data

Imaging data were processed using the SIMA^49^, Suite2p (https://www.biorxiv.org/content/10.1101/061507v2), and OASIS^50^ software packages. Motion correction was performed by concatenating all imaging sessions for a field of view and using the Suite2p rigid motion correction strategy. Only the GCaMP6f channel was used for estimating motion artifacts. Regions of interest (ROIs) were then drawn over putative CA1 pyramidal cell somata visible in the cross-session GCaMP6f channel time-average image. To prevent the introduction of any bias, the red mCherry channel time-average image was not viewed when drawing ROIs but only referenced after all ROIs had been drawn in order to tag ROIs over mcherry-expressing cells as Robo2 KO cells. In cases where no red channel was simultaneously recorded, an affine transformation was calculated to align the red-only image acquired just prior to imaging to the GCaMP6f channel time-average image.

Dynamic GCaMP6f fluorescence signals for each imaging session were extracted from the binary ROIs using SIMA. One session with dropped data during the last > 1 minute of recording was removed from analysis. Deconvolved spikes were computed for each ROI for each imaging session according to the following procedure: a baseline was calculated by smoothing the fluorescence trace with a Gaussian filter (std = 5 frames, window = 30s), calculating the rolling minimum of this smoothed trace, (window = 30s), and then calculating the rolling maximum of this last trace (window = 30). A preliminary spike train was estimated by deconvolving the baseline-subtracted trace with OASIS according to an AR1 model with ℓ_1_ penalty and pre-computed decay parameter using 400 ms for the GCaMP6f decay time. Using this spike train, a noise threshold was identified by computing the median and median absolute deviation (MAD) of the baseline-subtracted trace where no spikes were detected (i.e., putative noise-only time points). The noise threshold was defined as the median plus 3 MADs. A new spike train was then computed as before, now with the minimum spike size set explicitly to the noise threshold, and sparsity parameter, λ, set to 7.

#### Spatial Tuning Analysis

When evaluating the spatial tuning of PCs, we restricted our analysis to running-related epochs, defined as consecutive frames of forward locomotion at least 1 s in duration and with a minimum peak speed of 5 cm/sec. Consecutive epochs separated by <0.5 s were merged. Position was discretized into 100 2cm bins. Deconvolved spikes were smoothed with a Gaussian filter (std = 1 frame) and binarized. The spatial tuning vector was calculated as 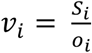, where *S_i_* is the number of running frames with spikes occurring at position i and *o_i_* is the number of running frames acquired at position i. In order to assess the significance of the spatial selectivity, for each cell we generated a null tuning distribution by cyclically permuting the position vector (restricted to running frames) by a random offset and repeatedly recomputing the tuning vector. This process was repeated 1,000 times. The true and null tuning vectors were then smoothed with a Gaussian (std = 3 position bins). Place fields were identified as at least 5 consecutive position bins above the 95^th^ percentile of the null distribution, in which the cell fired on at least 15% of laps.

#### Statistics

All tests are described in the appropriate figure legends. For imaging data, metrics were aggregated across all cells and sessions for a FOV, and FOVs were the unit of analysis for all statistical tests conducted. A paired student’s t test was used for comparison of means in paired data with n − 1 degrees of freedom. For the decoding analysis, a repeated measures two-way ANOVA with interaction was used to test the difference between subpopulation means across surrogate population sizes.

#### Code Availability

Suite2p and OASIS are both open source. The SIMA code base is open source and accessible at https://github.com/losonczylab/sima. All other code is available upon request, with a minimum working repository to be hosted publicly on github upon publication.

---

**Supplementary Figure 1 (related to Figure 2).**
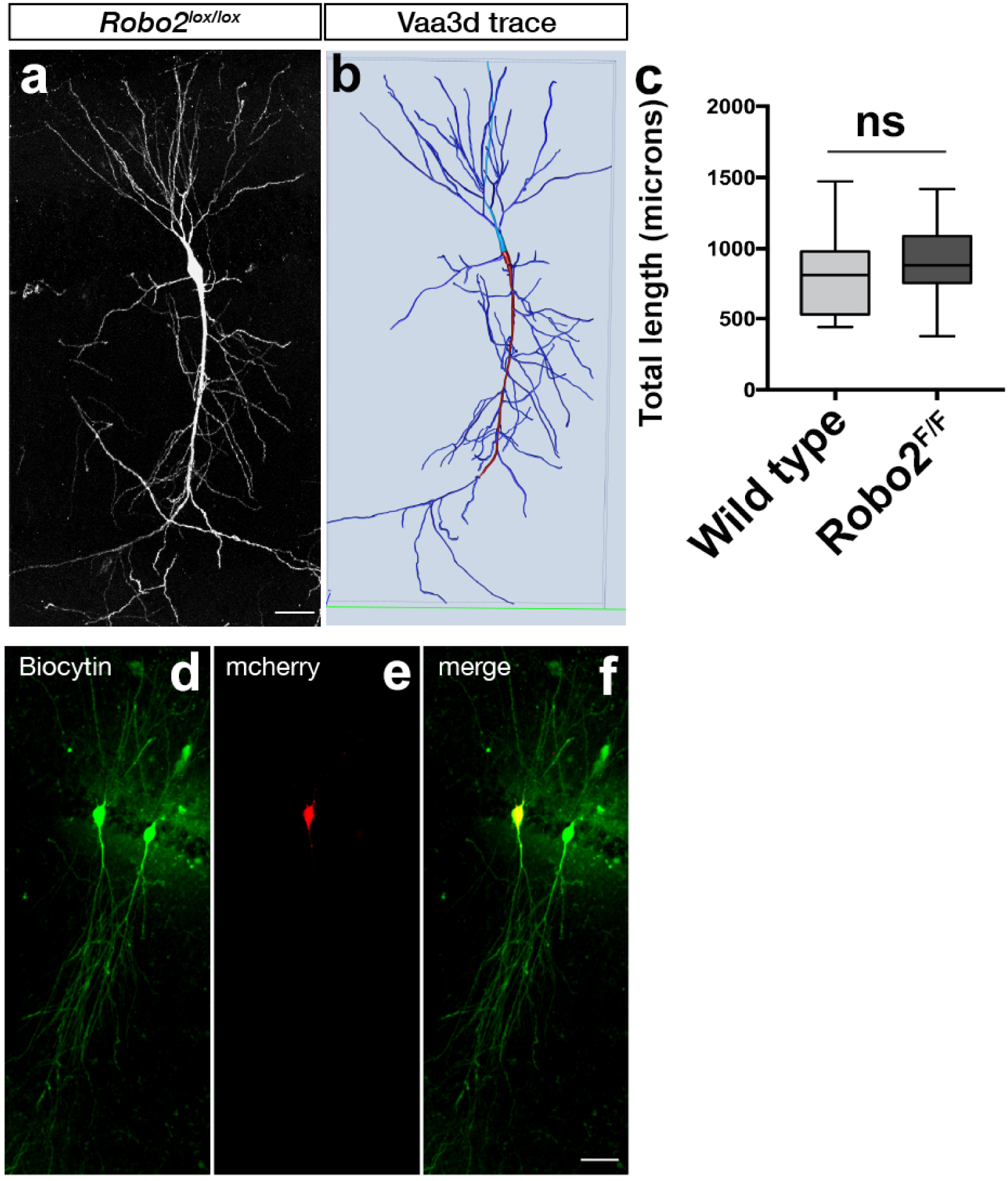
**a.** Robo2 is not required for dendrite growth. Dendritic path length analysis shows similar dendrite coverage between WT and Robo2 KO neurons. Neurons were reconstructed in 3D using the vaa3d platform (https://www.biorxiv.org/content/10.1101/395384v1). Scale bar: 100µm. **b.** Representative image of biocytin fill and *post-hoc* Streptavidin488 staining to unambiguously identify genotype and location of patched cells. Scale bar: 20µm.

**Supplementary Figure 2 (related to Figure 4).**
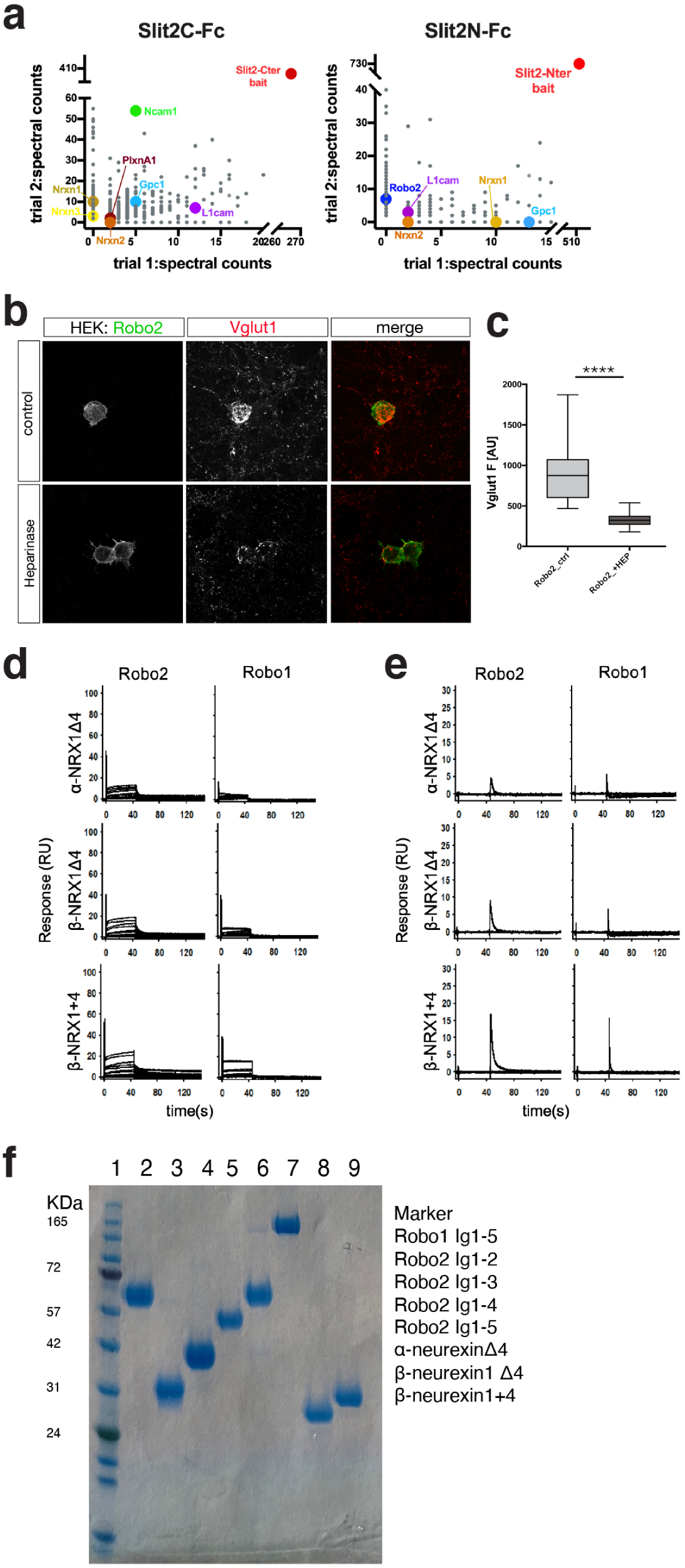
**a.** Mass-spectrometry results from synaptosome pulldown using SlitC-Fc and SlitN-Fc. Highlighted are cell surface adhesion proteins. Plotted data represents Ig-subtracted peptides only. **b.** Treatment of *in vitro* hemisynapse assay with Heparinase reduces Robo2-dependent Vglut1 clustering, scale bar: 7µm (****p<0.0001, Mann-Whitney). **c,d.** SPR binding experiments of Robos over NRXs: Binding of Robo2 and Robo1 (Ig1-5) over surfaces immobilized with α-NRX1Δ4, β-NRX1Δ4, and β-NRX1+4 ectodomains**. b.** Binding of Robos in the absence of heparin and **c.** binding traces of Robos in the presence of 3mM EGTA, instead of 3mM CaCl_2_. **e.** Coommassie gel showing quality and purity of all proteins used in SPR experiments.

